# The α-dystroglycan N-terminus is a broad-spectrum antiviral agent against SARS-CoV-2 and enveloped viruses

**DOI:** 10.1101/2023.11.06.565781

**Authors:** Maria Giulia Bigotti, Katja Klein, Esther S. Gan, Maria Anastasina, Simon Andersson, Olli Vapalahti, Pekka Katajisto, Maximilian Erdmann, Andrew D. Davidson, Sarah J. Butcher, Ian Collinson, Eng Eong Ooi, Giuseppe Balistreri, Andrea Brancaccio, Yohei Yamauchi

**Affiliations:** Bristol Heart Institute, Research Floor Level 7, Bristol Royal Infirmary; Bristol BS2 8HW, UK; School of Biochemistry, Faculty of Life Sciences, University of Bristol; Bristol BS8 1TD, UK; School of Cellular and Molecular Medicine, Faculty of Life Sciences, University of Bristol; Bristol BS8 1TD, UK; Program in Emerging Infectious Diseases, Duke-NUS Medical School; 8 College Road, Singapore, 169857, Singapore; Faculty of Biological and Environmental Sciences, Molecular and Integrative Biosciences Research Program, University of Helsinki; Helsinki, Finland; Helsinki Institute of Life Sciences-Institute of Biotechnology, University of Helsinki; Helsinki, Finland; Department of Virology, Medicum, Faculty of Medicine, University of Helsinki; Helsinki, Finland; Department of Virology, University of Helsinki and Helsinki University Hospital; Helsinki, Finland; Department of Biosciences and Nutrition, Karolinska Institutet; 141 83 Huddinge, Sweden; Department of Cell and Molecular Biology, Karolinska Institutet; 171 77 Solna, Sweden; Viral Research and Experimental Medicine Centre, SingHealth Duke-NUS Academic Medical Centre; 20 College Road, Singapore, 169856, Singapore; Saw Swee Hock School of Public Health, National University of Singapore; 12 Science Drive 2, #10-01, Singapore, 117549, Singapore; Institute of Chemical Sciences and Technologies “Giulio Natta” (SCITEC)-CNR; Rome, Italy; Institute of Pharmaceutical Sciences, Department of Chemistry and Applied Biosciences (D-CHAB), ETH Zurich; 8093 Zurich, Switzerland; Division of Biological Science, Graduate School of Science, Nagoya University; Furo-cho, Chikusa-ku, Nagoya, 464-8601, Japan

**Keywords:** extracellular matrix, α-dystroglycan, SARS-CoV-2, coronaviruses, enveloped viruses, broad-range antiviral

## Abstract

The COVID-19 pandemic has shown the need to develop effective therapeutics in preparedness for further epidemics of virus infections that pose a significant threat to human health. As a natural compound antiviral candidate, we focused on α-dystroglycan, a highly glycosylated basement membrane protein that links the extracellular matrix to the intracellular cytoskeleton. Here we show that the N-terminal fragment of α-dystroglycan (α-DGN), as produced in *E. coli* in the absence of post-translational modifications, blocks infection of SARS-CoV-2 in cell culture, human primary gut organoids and the lungs of transgenic mice expressing the human receptor angiotensin I-converting enzyme 2 (hACE2). Prophylactic and therapeutic administration of α-DGN reduced SARS-CoV-2 lung titres and protected the mice from respiratory symptoms and death. Recombinant α-DGN also blocked infection of a wide range of enveloped viruses including the four Dengue virus serotypes, influenza A virus, respiratory syncytial virus, tick-borne encephalitis virus, but not human adenovirus, a non-enveloped virus *in vitro*. This study establishes soluble recombinant α-DGN as a broad-band, natural compound candidate therapeutic against enveloped viruses.

## INTRODUCTION

Dystroglycan (DG) is an adhesion complex expressed in developing and adult tissues, where its main function is to anchor elements of the extracellular matrix (ECM) to the actin cytoskeleton and increase their stability. DG is present as part of the dystrophin-glycoprotein complex (**Fig. 1A, left**) in a variety of tissues and cell types, including skeletal and cardiac muscle, epithelial cells, neurons and glia in the central and peripheral nervous system (*1, 2*). Dystroglycan is composed of two subunits: the highly glycosylated extracellular α-subunit (α-DG) and the transmembrane β-subunit (β-DG), that interact non-covalently to form a bridge between the ECM and the actin cytoskeleton (**Fig. 1A, left**). α-DG is a receptor for ECM proteins such as laminins, perlecan, neurexins and agrin. It is composed of a N- and a C-terminal globular domain, separated by an elongated, extensively glycosylated mucin-like region, a domain arrangement evolutionarily remarkably conserved (*3*) (**Fig. 1A, left**). α-DG acts as the cell receptor for some arenaviruses such as Lassa fever virus (LFV) (*4*), that specifically interact with the glycans decorating the mucin-like region of α-DG (*5*). The proprotein convertase furin cleaves α-DG during maturation in the Golgi apparatus, liberating its N-terminal domain called α-DG N-terminus (α-DGN) (**Fig. 1A**, dubbed α-Nt in the dotted circle and zoomed on the right), during maturation in the Golgi apparatus (6). Contrary to the rest of the protein, α-DGN is scarcely glycosylated (*7*) and it is an autonomous folding unit that constitutes itself of an N-terminal Ig-like domain connected by a flexible loop to a ribosomal RNA binding protein S6-like domain (*8*) (**Fig. 1A, right**). Whilst the α-DGN domain harboured by α-DG has been shown to be important for α-DG decoration with glycans (*4, 9*), the physiological function of the α-DGN is unknown, despite its detection in trace amounts in plasma and cerebrospinal fluids (*10–12*). It was recently reported that α-DGN exerts antiviral activity against Influenza A virus in mice (*13*). Here, we show that α-DGN is a broad range antiviral protein that blocks infection by a wide range of enveloped viruses including severe acute respiratory syndrome coronavirus 2 (SARS-CoV-2) and dengue virus.

**Fig. 1.**
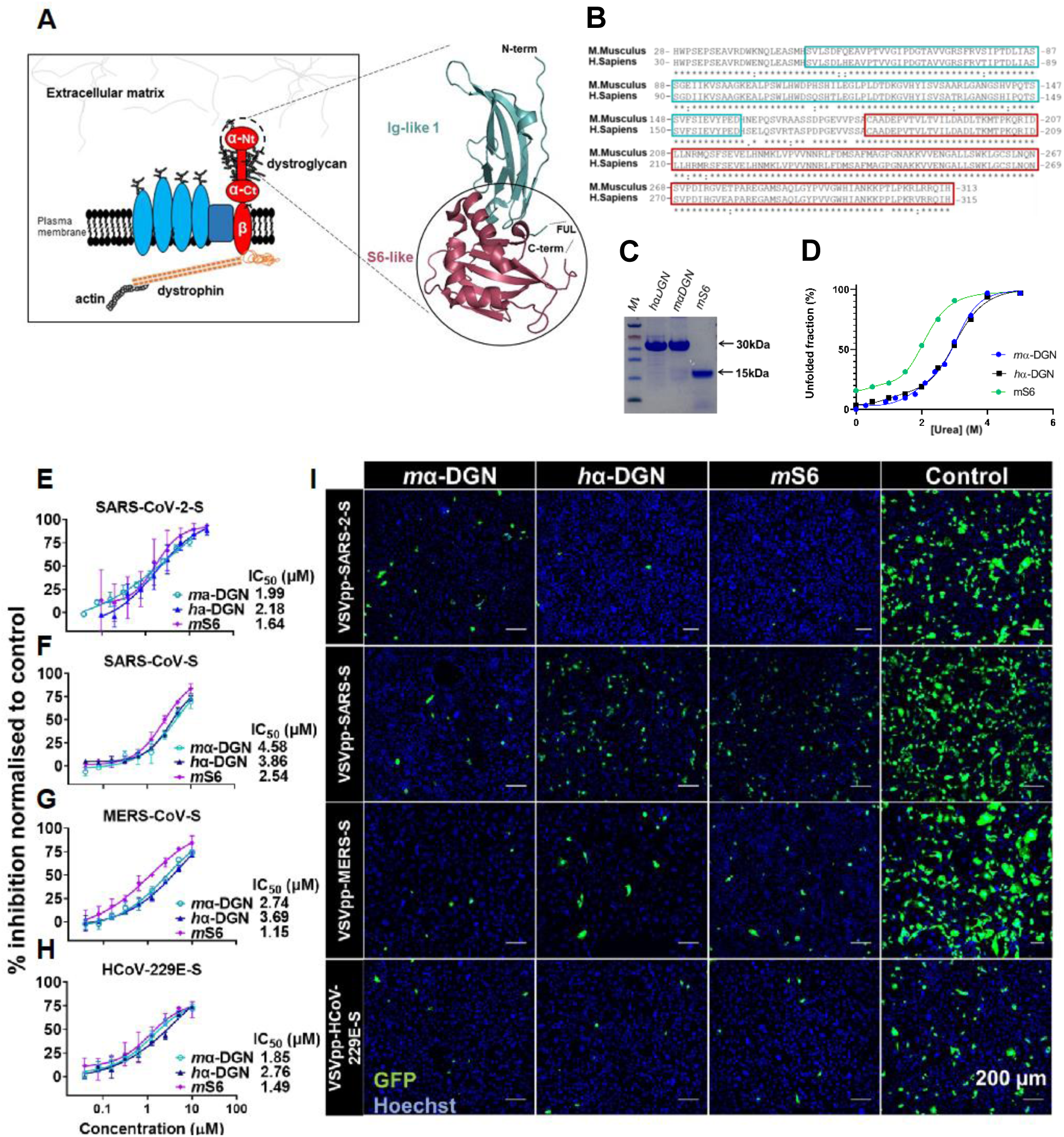
α-DGN and its inhibitory activity against pseudotyped human coronaviruses. (**A**) **Left**: schematic representation of the dystrophin-glycoprotein complex (DGC). This multiprotein complex, of which dystroglycan (DG) is a central element, anchors the extracellular matrix (ECM) to actin and other components of the cytoskeleton. Extracellular α-DG binds different ECM proteins, such as laminin, while transmembrane β-DG binds the actin cytoskeleton *via* direct interaction with dystrophin. Also depicted are other intracellular molecules associated with the DGC. The N-terminal of α-DG (α-DGN), highlighted in the dashed circle, is liberated in circulation following cleavage by furin. α-Ct: α-DG C-terminal, α-Nt: α-DG N-terminal. **Right**: cartoon representation of the 3D-structure of α-DGN (PDB: 1U2C). Shown are the two domains (N-term Ig-like, in cyan and C-term S6-like, in magenta and enclosed into the full circle) and the position of the flexible undefined loop (FUL) connecting them is indicated. (**B**) Alignment of the amino acid sequences of *m*α-DGN (Uniprot: Q62165, top) and *h*α-DGN (Uniprot: Q14118, bottom), showing a degree of identity of ∼93 % (∼97 % similarity). The sequences of the Ig-like and S6-like domains are highlighted in cyan and magenta boxes, respectively. Amino acid conservation code: (*) identity, (:) strong similarity, (.) weak similarity. (**C**) SDS-PAGE of the recombinant α-DGN proteins (indicated over each lane) used in this study, as final products of the purification procedure. (**D**) Recombinant α-DGN unfolding curves measured by intrinsic tryptophan fluorescence spectroscopy, as fitted to a two-state linear extrapolation model. The mid-point of the transitions (Cm), 2.8M for *m*α-DGN, 2.7M for *h*α-DGN and 1.8M for the shortened version *m*S6, are indicative of stable polypeptides. (**E-H**) Dose-response curves for *m*α-DGN, *h*α-DGN or *m*S6 pre-treated Caco-2 cells infected with VSV pseudotyped particles expressing the Spike protein from E) SARS-CoV-2, F) SARS-CoV, G) MERS-CoV and H) HCoV-229E. Infection was quantified by measuring GFP-positive cells and data are expressed as % inhibition normalized to an unrelated protein control (mean ± SEM). IC_50_ values (from n=3 independent experiments) were calculated using nonlinear regression calculations. (**I**) Representative immunofluorescence images of infected cells treated with either *m*α-DGN, *h*α-DGN, *m*S6 or vehicle control (10 µM), as indicated (infected, GFP-positive cells in green and Hoechst for nuclei in blue). Scale bar: 200 µm.

## RESULTS

### α-DGN and its S6-like domain block pseudotyped human coronavirus infection

We used *E. coli* to produce recombinant α-DGN from *M. musculus* (*m*α-DGN, residues 50-313) and *H. sapiens* (*h*α-DGN, residues 52-315) (**Fig. 1B)** in the pHis-Trx expression vector *14* either in its original form or modified as described in the Materials and Methods. The proteins were stabilised by replacing an Arg residue located in the loop region connecting the Ig-and S6-like domains (R166 in *m*α-DGN and R168 in *h*α-DGN) (*8*) with a His residue **(Fig. 1A, right** and **Fig. S1A)**. Both, α-DGN proteins, which share 92.7 % sequence identity **(Fig. 1B**), ran as homogeneously pure electrophoretic bands at an apparent molecular weight of 30 kDa (**Fig. 1C)**. The homogeneity of the protein preparations was further confirmed by electrospray ionization liquid chromatography mass spectrometry (**Fig. S1B**). The proteins were stable as measured by intrinsic tryptophan fluorescence upon urea-mediated denaturation (**Fig. 1D**).

We tested the antiviral activity of *m*α-DGN and *h*α-DGN against infection of Caco-2 cells by vesicular stomatitis virus (VSV) pseudotyped with the spike (S) proteins of human coronaviruses (CoVs) SARS-CoV-2, SARS-CoV, Middle East respiratory syndrome coronavirus (MERS-CoV) and human coronavirus 229E (HCoV-229E). Both *m*α-DGN and *h*α-DGN had potent antiviral activity against all pseudoviruses tested. The IC_50_ values for SARS-CoV-2-S, SARS-CoV-S, MERS-CoV-S and HCoV-229E-S were 1.99, 4.58, 2.74, and 1.85 µM for *m*α-DGN, and 2.18, 3.86, 3.69, and 2.76 µM for *h*α-DGN, respectively **(Fig. 1E-H and I).** The ECM protein agrin, recombinantly expressed from *E. coli*, was used as a non-specific protein control. These results established that α-DGN has broad antiviral activity against VSV pseudotyped with different coronavirus spikes.

Based on our previous experience in producing shortened versions of *m*α-DGN [8], we then dissected it into its two main domains, the Ig-like and the S6-like domains, to analyse their possible inhibitory potency. We cloned them separately into the pHisTrx(TEV) vector and produced each as described for the full-length protein. The Ig-like domain could not be purified efficiently due its low stability. However, the purified S6-like domain (hereafter *m*S6) was stable in solution (**Fig. 1D**) as a ∼15 KDa protein and was further tested for its antiviral activity as described above. *m*S6 inhibited infection of VSV pseudotyped with CoV-S proteins from SARS-CoV-2, SARS-CoV, MERS-CoV and HCoV-229E with IC_50_ values of 1.64, 2.54, 1.15 and 1.49 µM respectively (**Fig. 1E-H**). This showed that *m*S6 blocks CoV-S-pseudotyped VSV infection with comparable potency to the full-length *m*α-DGN.

### α-DGN blocks different SARS-CoV-2 variant infection in susceptible cell lines

In the first instance, we tested *m*α-DGN against the clinical ancestral SARS-CoV-2 isolate Finland/1/2020 (in HEK 293T cells stably expressing human angiotensin-converting enzyme 2 (hACE2) and human transmembrane serine protease 2 (TMPRSS2) (*15, 16*). Here, *m*α-DGN blocked SARS-CoV-2 infection in a dose-dependent manner, with a 87.2 % infection reduction compared to the buffer control (20mM TrisHCl, 100mM NaCl, pH 7.5) at the highest concentration tested (10 µM) **(Fig. S2A)**. We further tested the inhibitory activity of *m*α-DGN against infection of VeroE6-TMPRSS2 (VTN) cells with different SARS-CoV-2 variants. The tested variants included the ancestral strain and four variants of concern (VOCs) including Gamma, Delta, Omicron BA.1 and Omicron BA.4 that evolved during the SARS-CoV-2 pandemic. Pre-treatment of cells with 10 µM *m*α-DGN blocked infection of VeroE6-TMPRSS2 cells with SARS-CoV-2 ancestral strain, Gamma, Delta, Omicron BA.1 and Omicron BA.4 by 80.4, 61.7, 86.4, 84.3 and 82.7 % respectively (**Fig. 2A and B**). These data further support the broad-band antiviral activity of α-DGN against authentic SARS-CoV-2 VOCs.

**Fig. 2.**
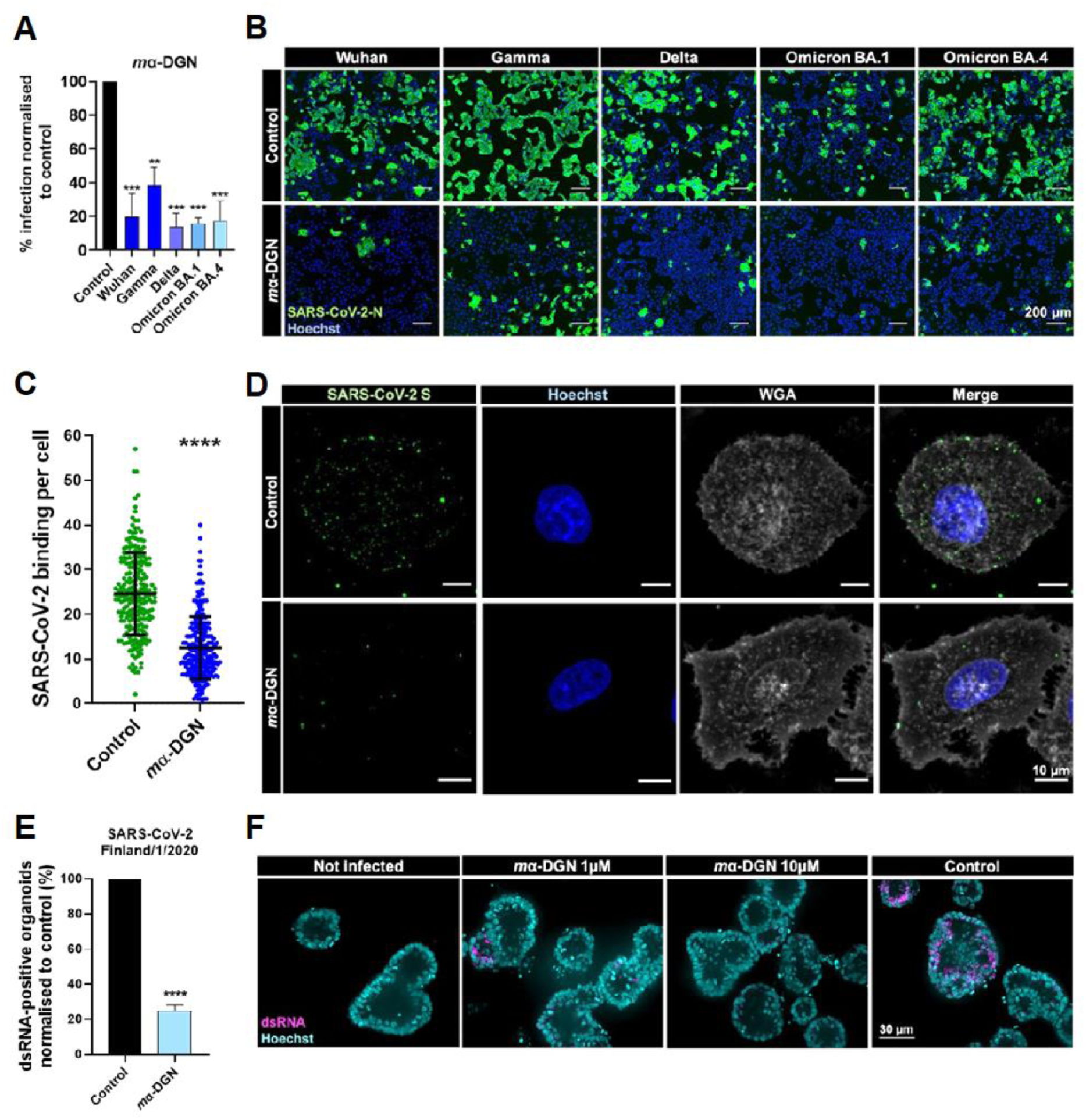
α-DGN blocks SARS-CoV-2 in epithelial cells and human primary gut organoids. (**A**) α-DGN inhibits infection of cells from different SARS-CoV-2 variants. VeroE6-TMPRSS2 cells were pre-treated with 10 µM *m*α-DGN before infection with SARS-CoV-2 Wuhan, Gamma, Delta, Omicron BA.1 or Omicron BA.2 (MOI=0.1). Infection was quantified using viral N protein staining (n=3 independent experiments). Bars are means + SEM. Scale bar, 200 µm. p-values were determined by one-way ANOVA with multiple comparison with respect to the control. p***<0.001, p**<0.01. (**B**) Representative immunostaining images of control or *m*α-DGN treated VeroE6-TMPRSS2 cells infected with the indicated variants of SARS-CoV-2 (viral N protein in green and Hoechst for nuclei in blue). (**C**) *m*α-DGN inhibits SARS-CoV-2 binding to cells. HeLa-ACE2 cells were pre-incubated with 10 µM *m*α-DGN or vehicle control, after which SARS-CoV-2 was added for 60 min on ice, and cells were fixed. SARS-CoV-2 binding per cell after treatment with *m*α-DGN (blue) or control (green) was quantified for >100 cells per condition. Significance was determined using a Mann-Whitney test (n=3 independent experiments), p****<0.0001. (**D**) Representative immunostaining images of extracellular SARS-CoV-2 particles on HeLa-ACE2 cells after pre-incubation with vehicle control or *m*α-DGN. Cells were stained with Hoechst for nuclei (blue), wheat germ agglutinin (WGA) for cell outline (grey) and for SARS-S protein (green). Scale bars, 10 µm. (**E**) *m*α-DGN blocks SARS-CoV-2 infection of human primary gut organoids. Organoids were treated with 1 or 10 µM of *m*α-DGN or buffer control and infected with SARS-CoV-2 at MOI 1 for 48 h before fixation. The graph shows the percentage of infected (double stranded RNA-positive) organoids, as normalised to the buffer treated control (mean + SD, n=4). (**F**) Representative immunostaining images of human gut organoids treated with indicated concentrations of *m*α-DGN or vehicle (PBS) control and infected with SARS-CoV-2 (dsRNA in magenta and nuclei are stained with Hoechst in cyan). Images represent the max Z-projection of 5 optical slices (out of 180 for each image stack) showing the mid-section of the organoids. Scale bars, 30 μm.

### α-DGN acts at the early stages of viral infection

To test what stage of viral infection is targeted by *m*α-DGN, we analysed its inhibitory activity on the virus as a function of time of administration. Pre-incubation of SARS-CoV-2 with *m*α-DGN before addition to the cells resulted in 50 % inhibition, while adding *m*α-DGN and SARS-CoV-2 at the same time to the cells reduced infection by 76.4 %. In contrast, addition of *m*α-DGN 2 h after SARS-CoV-2 inoculation resulted in little or no reduction of infection (**Fig. S2B**). We tested whether *m*α-DGN affects intracellular SARS-CoV-2 replication using the pSC2-Rep-Wu-Gp-RL replicon that lacks the S and M gene coding sequences (preventing viral assembly and egress) (*17*). RNA replication of the replicon can be measured by the encoded Renilla luciferase (RLuc). Immediately after electroporation with the replicon RNA transcripts, VTN cells were treated with 10 µM *m*α-DGN, buffer control, or remdesivir, a known inhibitor of the viral RNA-dependent RNA polymerase synthesis. Upon incubation for 18 h, we observed no difference in RLuc activity between the mα-DGN-and buffer control-treated cells (**Fig. S2C**) or the immunofluorescence staining of the dsRNA (a marker for viral/replicon RNA replication) (**Fig. S2D and E**). This confirmed that α-DGN does not block intracellular viral genome replication, in contrast to remdesivir.

When HeLa cells expressing hACE2 were pre-incubated with *m*α-DGN before virus addition, SARS-CoV-2 binding was blocked by 50 % compared to the buffer control (**Figure 2C and D**). Incubation of AAT or Caco-2-ACE2 cells with 4 µM Alexa Fluor-648 labelled *m*α-DGN in the cold for 30 min revealed that α-DGN binds to the cell surface (**Fig. S3A and B**). α-DGN binding was observed in both AAT and Caco-2-ACE2 cells, suggesting that α-DGN cell binding is not cell-type specific.

### α-DGN blocks SARS-CoV-2 in human primary gut organoids

Organoids are miniaturized three-dimensional organs assembled *in vitro* with stem cells or adult cells from specific tissue types. In COVID-19 patients, SARS-CoV-2 infects the digestive tract and has been detected in the small intestine, in the colon and in the ileum (*18*). Therefore, human primary gut organoids are employed as a model system in SARS-CoV-2 studies for their superior capacity to mimic the complexity of human tissue and organs in physiological as well as pathological conditions, as compared to 2D cell culture models (*18*). To analyse the antiviral activity of *m*α-DGN in these organoids, we first amplified duodenum tissue explants of human donors *in vitro* as three-dimensional multi-crestal organoids. After mechanical disruption to expose the cellular apical surface, the gut organoids were used for inhibition studies. Pre-treatment of SARS-CoV-2 isolate Finland/1/2020 with 10 µM *m*α-DGN decreased the number of infected organoids by 74.9 % (from 20.7 % in the vehicle control to 5.2 % in *m*α-DGN-treated organoids) **(Fig. 2E and F)**. This shows that *m*α-DGN efficiently inhibits SARS-CoV-2 infection in in an organoid model.

### α-DGN inhibits SARS-CoV-2 in humanised ACE2 mice

We further examined the therapeutic and prophylactic potential of αDGN *in vivo* using the SARS-CoV-2 infection model in K18-hACE2 transgenic mice expressing hACE2 (*19*). First, mice were treated with a single dose of 7.5 or 0.75 µg of *m*α-DGN per mouse via the intranasal (IN) route immediately followed by an IN lethal dose (5×10^4^ TCID_50_) of SARS-CoV-2 (SG12-B strain (*19*)). All animals were monitored daily for weight loss and survival (**Fig. 3A**). We observed that after one IN administration of α-DGN, mice were protected from a lethal SARS-CoV-2 infection, with a survival rate increase of 20 % and delayed weight loss (**Fig. 3B and C**). Moreover, treated mice had significantly reduced endpoint viral loads in their lungs compared to animals receiving only a buffer control (**Fig. 3D**). We then determined the prophylactic potential by administering *m*α-DGN or *h*α-DGN at 7.5 µg/mouse/dose via the IN route two hours before SARS-CoV-2 (SG12-L strain (*19*)) challenge, followed by one daily IN administration of the same *m*α-DGN or *h*α-DGN dose for the next three consecutive days (**Fig. 3E**). Animals treated with *m*α-DGN or *h*α-DGN showed increased time of survival (**Fig. 3F**) without detectable weight loss delay (**Fig. 3G**) as well as significantly reduced lung viral loads (**Fig. 3H**) compared with control mice. Taken together, these studies demonstrate that αDGN increased survival and reduced respiratory infection in humanised ACE2 mice upon challenge with a lethal dose of SARS-CoV-2.

**Fig. 3.**
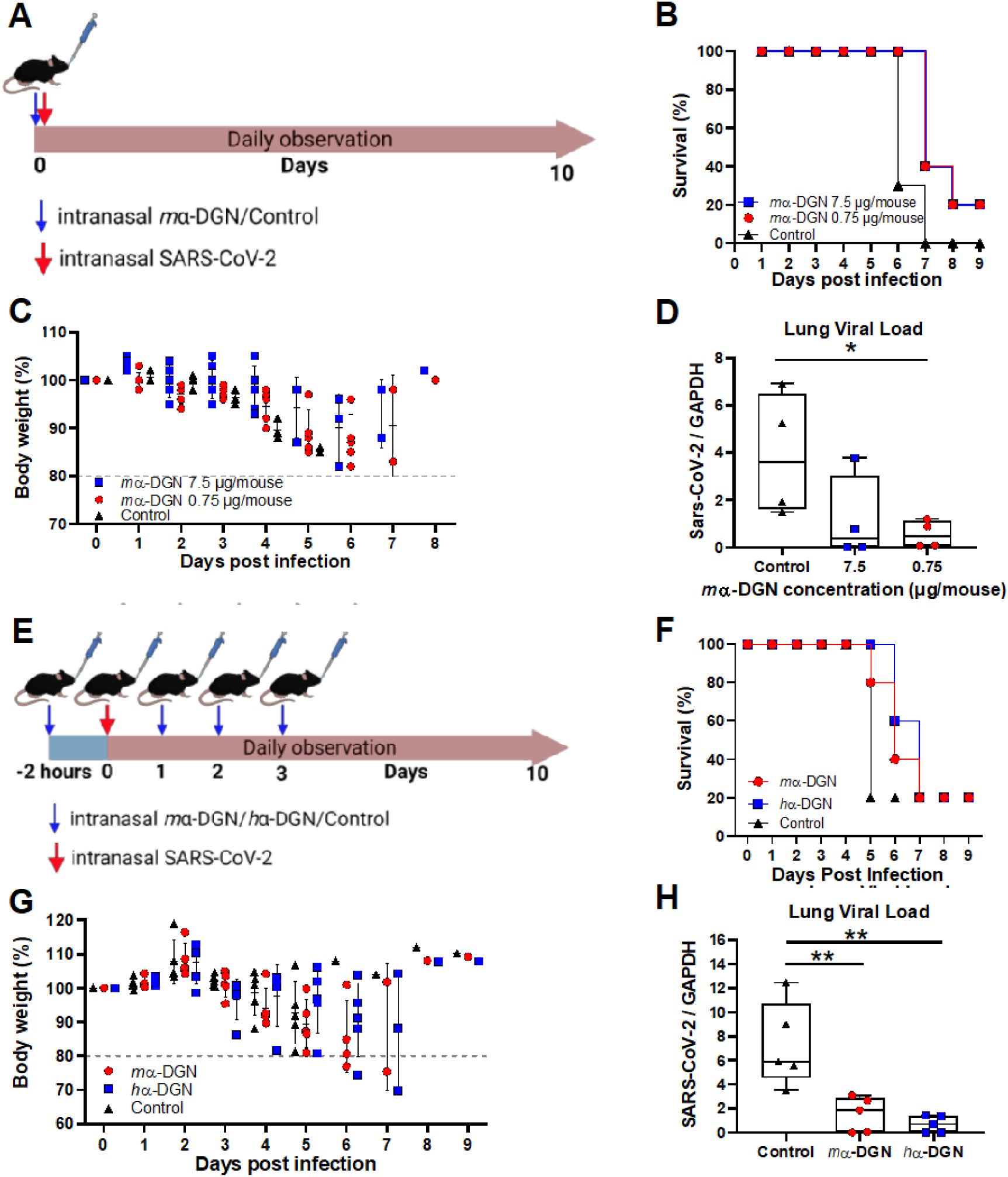
αDGN inhibits SARS-CoV-2 infection in K18-hACE2 mice. (**A**) *m*α-DGN or buffer control was administered to K18-hACE2 mice at 7.5 µg/mouse or 0.75 µg/mouse at the same time as the lethal dose of the challenging virus (SARS-CoV-2 SG12-B) via the intranasal (IN) route. Mice were observed daily for (**B**) survival (n=5 per group) and (**C**) weight loss (n=5 per group). (**D**) Lung viral loads as determined at end point (n=4 per group). Statistical significance was determined using a Mann Whitney test, p*<0.05. ns: not significant. (**E**) Buffer control, *m*α-DGN or *h*α-DGN at 7.5 µg/mouse were administered via the IN route 2 h prior to virus challenge (SARS-CoV-2 SG12-L) and then daily for 3 days. Mice were then observed daily for (**F**) survival (n=5 per group) and (**G**) weight loss (n=5 per group). (**H**) Lung viral loads as determined at endpoint (n=5 per group). Statistical significance was determined using a Mann Whitney test, p** <0.01.

### α-DGN blocks a broad range of enveloped viruses

The measured inhibitory effect of α-DGN on the infectivity of pseudotyped CoVs and SARS-CoV-2 variants prompted us to investigate its broad antiviral potential against other human viruses. Specifically, we further tested *m*α-DGN for its antiviral activity against other enveloped viruses: influenza A virus (IAV) X-31 (H3N2), respiratory syncytia virus (RSV), Semliki Forest virus (SFV), tick-borne encephalitis virus Kuutsalo-14 isolate (TBEV), vesicular stomatitis virus (VSV) and Dengue virus (DENV), and the non-enveloped human adenovirus serotype 5 (hAdV5). Infection assays were performed in a multiplexed format and automated confocal fluorescence microscopy followed by image analysis was used to determine the fraction of infected cells. Infected cells were identified by immunostaining using antibodies specific for the IAV viral nucleoprotein, TBEV membrane protein (*20*) or by detecting the green or red fluorescence of fluorescent reporter genes inserted in the genome of each virus by reverse genetic engineering, specifically: GFP for SFV (*21*), RSV (*22*) and VSV (*23*), and RFP for hAd5, the latter kindly provided by Dr Vincenzo Cerullo, University of Helsinki. *m*α-DGN (10 µM) reduced infection of all enveloped viruses i.e., IAV by 80 %, RSV by 91 %, SFV by 57 %, TBEV by 90 %, VSV by 99 % (**Fig. 4A-E**). *m*α-DGN reduced the replication of Dengue virus (DENV) serotypes 1, 2, 3 and 4 by 57, 58, 50 and 42.5 %, respectively, as detected by qRT-PCR **(Fig. 4G-J).** *m*α-DGN also blocked DENV4 infection of human primary monocytes by 91.3 % (**Fig. S4**). *m*α-DGN did not inhibit the non-enveloped virus hAdV5 (**Fig. 4F**). Our observations suggest that α-DGN is a broad-range inhibitor against enveloped viruses.

**Fig. 4.**
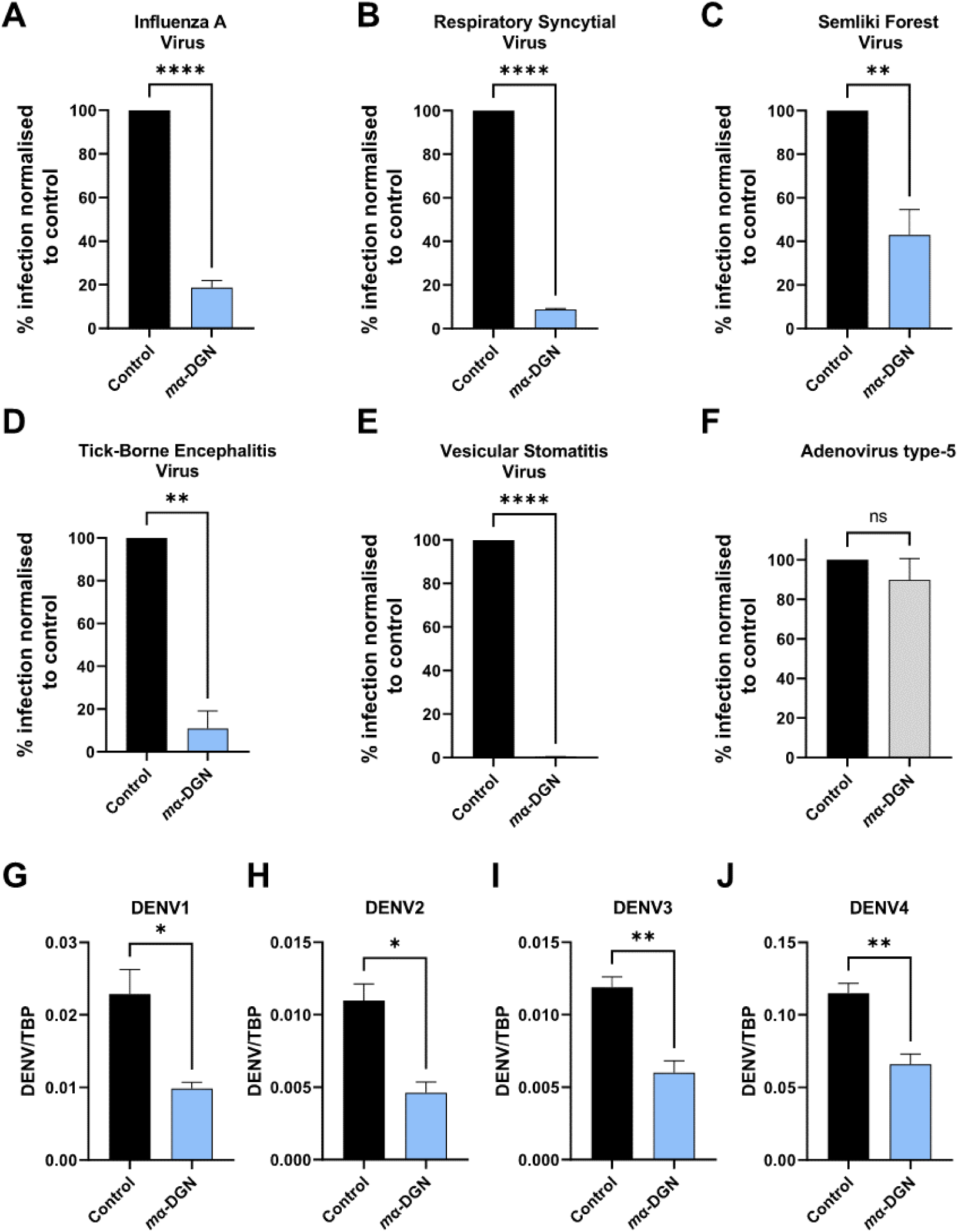
*m*αDGN inhibits a broad range of enveloped viruses. (**A-F**) Inhibitory activity of *m*α-DGN in infection assays against (**A**) IAV, (**B**) RSV, (**C**) SFV, (**D**) TBEV, (**E**) VSV and (**F**) AdV5. Infection was analysed by immunostaining and results are represented as % inhibition normalised to the untreated control. (**G-J**) Inhibitory activity of *m*α-DGN against the 4 Dengue Virus serotypes (**G**) DENV1, (**H**) DENV2, (**I**) DENV3 and (**J**) DENV4 in cell assays. DENV RNA was analysed by qRT-PCR. DENV copies were normalised to the internal control protein TBP. Values are means + SD. Statistical significance was determined using an unpaired t-test. ns = not significant, p* < 0.1, p** <0.01, p*** <0.001.

## DISCUSSION

The first evidence of the involvement of the extracellular matrix complex dystroglycan in viral infection came with the identification of α-DG as the cellular receptor for arenaviruses such as Lassa fever virus (LFV) and lymphocytic choriomeningitis virus (LCMV) (*4*). The interaction of α-DG with arenaviruses has been found to be specific for the Old World and clade C New World arenaviruses only (*24*), and to be dependent on the glycosylation shell of the so-called mucin-like region of α-DG (*25*). The mucin-like region is comprised of the amino acids linking the C-terminal and N-terminal globular domains of α-DG (**Fig. 1A, left**). It is highly decorated with long, linear chains of repeating carbohydrate units collectively called *matriglycan*, whose recognition by LASV has been recently characterized in detail (*5*) and demonstrated to be key in viral entry. In our study, we used a recombinant non-glycosylated N-terminal globular domain of α-DG (α-DGN) produced in *E. coli*. Therefore, the antiviral mechanism is independent of its glycosylation.

The emergence of the COVID-19 pandemic prompted us to explore the potential of α-DGN as a SARS-CoV-2 inhibitor. de Greef and colleagues previously showed that transgenic mice lacking the *Dag1* gene (which encodes α-DG and β-DG) were more susceptible to IAV PR8 (H1N1) lung infection than control mice (*13*). Recombinant α-DGN inhibited IAV HA-mediated hemagglutination of chicken red blood cells (*13*), probably by blocking IAV binding to the red blood cell surface, similar to the inhibition of virus entry for SARS-CoV-2 (**Fig. 2D**). Adenoviral overexpression of α-DGN or administration of recombinant α-DGN decreased IAV PR8 titres in the lungs of mice (*13*).

Our extensive testing proved that our recombinant, stabilized form of α-DGN produced in *E. coli* blocked VSV pseudotyped with CoV S proteins including SARS-CoV-2, SARS-CoV, MERS-CoV and HCoV-229E and a diverse range of enveloped RNA viruses *in vitro* and *in vivo*. All the SARS-CoV-2 VOCs tested were blocked by α-DGN, as well as enveloped viruses that cause respiratory disease (IAV, RSV), fever (SFV, DENV), viral meningitis (TBEV) in humans and lesions in livestock (VSV) (**Fig. 2 and 4**). Importantly, the blockade of all four DENV serotypes is particularly promising in the face of the difficulties in treating severe secondary infections caused by antibody-dependent enhancement (ADE) of different serotypes (*26*). α-DGN also delayed mortality and reduced lung viral loads both in a therapeutic and prophylactic administration regime of SARS-CoV-2 lethal challenge in K18-hACE2 mice **(Fig. 3)**. Our results show α-DGN to be highly effective when administered intranasally, which is the preferred route of administration of a protein-based therapeutic. Below are the potential advantages when considering α-DGN as a candidate antiviral compound:

i. **Facile production.** Recombinant α-DGN is inexpensive to produce, both the murine and human forms are stable in solution at ambient temperature and can be easily scaled up for mass production.
ii. **Discovery of the 15 KDa active domain S6.** The smaller size of *m*S6 compared with *m*α-DGN can be an advantage for translational research.
iii. **Weak immunogenicity potential.** α-DGN is produced by the host cell as a polypeptide. This feature, together with the very high degree of amino acid sequence conservation amongst species and the absence of post-translational modifications, makes α-DGN more unlikely to elicit an immune response when used as a therapeutic (*27*). Our recombinant *mα*-DGN and *h*α-DGN, produced in *E. coli* devoid of potentially immunogenic post-translational modifications, did not elicit any adverse response when administered to human cell lines, human gut organoids or mice.
iv. **a-DGN has comparable or superior IC_50_ to other antiviral compounds.** *m*α-DGN, *h*α-DGN and *m*S6 display potent broad-range antiviral activity. The IC_50_ values were in the low micromolar range for all the viruses tested. These values are higher than those of viral spike neutralizing monoclonal antibodies (nmAbs) that display IC_50_ values in the low nanomolar range. However, mAb-based therapies lead to lack of broad-spectrum activity as is evidenced by the altered efficacy of the same mAb against emerging variants of SARS-CoV-2 (*28*). The antiviral efficacy of α-DGN is comparable to that of small antiviral compounds currently in use against SARS-CoV-2 infection, such as remdesivir, molnupiravir or nirmatrelvir, whose IC_50_ values are also in the low micromolar range (*29*). Picolinic acid, a natural compound with a broad-band antiviral effect against enveloped viruses exhibits an IC_50_ of 0.5 mM against IAV PR8 in cell culture (*30*), which is approximately 200-fold higher than the IC_50_ values of α-DGN against CoVs.
v. **Antiviral efficacy is observed *in vivo* at low dosage**. Our *in vivo* studies using K18-hACE2 mice resulted in a significant reduction of the SARS-CoV-2 lung viral loads in a lethal challenge model. The daily dose of α-DGN in these mice was 0.3 mg/kg (single dose therapeutic treatments, and once a day for prophylactic treatments), which is 1333- and 2000-fold lower than that of molnupiravir (200 mg/kg, twice a day) and nirmatrelvir (300 mg/kg, twice a day) employed in other studies via the IN route (*31*).

In summary, we showed that α-DGN blocks SARS-CoV-2 infection at the step of cell binding, which may inhibit efficient cell-to-cell spread of the virus in lungs. Whether inhibition depends on binding of α-DGN to the host cell, to the virus, or both needs further investigation. Understanding the molecular mechanism of action of α-DGN antiviral activity will help elucidate the physiological relevance of trace α-DGN levels detected in human plasma (*10, 12*).

### Conclusions

We have characterized α-DGN, a naturally occurring protein, as a potent broad antiviral compound that blocks infection of SARS-CoV-2 in human cell lines, human gut organoid models and mouse models of infection. The antiviral activity of α-DGN is wide-spread among enveloped viruses of clinical impact that cause lethal human infections such as IAV, RSV and DENV. We conclude that α-DGN and its active domain S6-like is a promising lead compound for future research and development into a pan-enveloped virus antiviral.

## MATERIALS AND METHODS

### Study design

Our research aimed to test α-DGN as a potential broad-range antiviral protein to block infection of enveloped viruses *in vitro* and *in vivo*. We investigated’ antiviral activity of α-DGN against a wide range of enveloped viruses, including different coronavirus strains and variants, IAV, RSV, SFV, TBEV, VSV and DENV. Viral inhibition experiments were conducted with virus-corresponding susceptible cell lines or human primary cells and were performed in at least three independent biological replicates. We further characterised the time-dependent antiviral activity of α-DGN by time of addition, viral binding inhibition and intracellular replication studies in different cell lines. *In vivo* SARS-CoV-2 inhibition studies were performed in K18-hACE2 C57/BL/6 mice (n=5 mice per group). All *in vivo* experiments were performed according to the approved protocol by the Institutional Animal Care and Use Committee. Animals were randomly assigned to groups, but treatment was not blinded to the investigator. Sample size was determined based on previous studies.

### α-DGN production

The murine and human proteins (variants R166H and R168H, respectively) were produced in *E.coli* BL21 CodonPlus(DE3) RIL (Agilent Technologies) carrying the plasmid pHisTrx-mα-DGN and pHisTrx-hα-DGN, respectively, as previously described (*32*). Both plasmids were used either in their original form or in a mutated form where the thrombin cleavage site between the thioredoxin fusion protein and α-DGN was replaced by a TEV proteolytic site, to yield pHisTrx(TEV)-mα-DGN and pHisTrx(TEV)-hα-DGN. A multi-sites mutagenesis protocol adapted from the Quickchange single-step mutagenesis protocol was employed successfully to insert this modification (*33*). The gene of the mouse S6-like domain (for the amino acid sequence see fig S1A), as synthetically produced by GeneArt, was cloned into the pHis-Trx(TEV) plasmid for protein production in *E.coli* downstream a His-tagged thioredoxin fusion partner. The *m*S6 protein was expressed and purified following the same protocol as the full-length protein.

### Cell lines and antibodies

African Green Monkey kidney epithelial cell line Vero E6 expressing TMPRSS2 (VeroE6-TMPRSS2), human embryonic epithelial cells 293T expressing ACE2 and TMPRSS2 (HEK-Ace2-TMPRSS2) (*15*), human cervical epithelial cell line HeLa and HeLa cells expressing ACE2, human lung epithelial cell line A549 and A549 cells expressing ACE2 and TMPRSS2 (AAT) were cultured in Dulbecco’s Modified Eagle’s Medium with 4500 mg/L glucose (Sigma-Aldrich, FIN) supplemented with 10 % fetal bovine serum (FBS; Gibco, FIN, UK), 2 mM L-glutamine (Sigma-Aldrich, FIN, UK), 100 units of penicillin and 0.1 mg/ml streptomycin (PenStrep; Sigma-Aldrich, Fin, UK), and non-essential amino acids (NEAA; Sigma-Aldrich, FIN, UK). Human neuroblastoma cell line SK-N-SH (ATCC HTB-11) was cultured in Dulbecco’s Modified Eagle’s Medium with 1g/L glucose (Sigma-Aldrich, FIN) supplied with 10 % FBS, L-glutamine and PenStrep. The colorectal adenocarcinoma Caco-2 cell line and Caco-2 expressing ACE2 were maintained in DMEM+GlutaMAX containing 1mM Sodium Pyruvate (Gibco^TM^, Thermo Fisher, UK), 10% FBS (Gibco^TM^, Thermo Fisher, UK), PenStrep, and 0.1mM NEAA (Sigma Aldrich, UK). Huh7 (Duke Cell Repository) cells were cultured in Dulbecco’s Modified Eagle Medium (DMEM) with 10% FBS. All cell lines were cultured in a 37°C incubator with 5 % CO_2_ and passaged 1:8 (VeroE6-TMPRSS2, HeLa, HeLa-ACE2, A549, AAT), 1:5 (SK-N-SH) or 1:4 (Caco-2, Caco-2-ACE2) every three to four days.

Immunostaining was done using polyclonal rabbit anti-SARS-CoV Nucleocapsid (N) protein antibody (Rockland, UK), polyclonal rabbit anti-SARS-CoV-2 (2019-nCoV) Spike antibody (SinoBiological, UK), in house prepared monoclonal mouse hybridoma anti-NP (HB-65) antibody, rabbit polyclonal antibody commercially raised against TBEV M, goat anti-rabbit Alexa Fluor-488 (Thermo Fisher, UK), goat anti-mouse Alexa Fluor-488 (Thermo Fisher, UK), goat anti-rabbit Alexa Fluor-594 (Thermo Fisher, UK), Wheat Germ Agglutinin (WGA) Alexa Fluor-647 (Thermo Fisher, UK), phalloidin Alexa Fluor-594 (Thermo Fisher, UK), Hoechst 3342 (Thermo Fisher, UK).

### Viruses

The following SARS-CoV-2 ancestral early passage virus isolates and VOCs were used in this study: ancestral SARS-CoV-2 isolates Finland/1/2020 (GenBank ID: MT020781.2), SARS-CoV-2/hCoV-19/Singapore/2/2020 (SG12) (GISAID ID: EPI_ISL_407987) (*19*), SARS-CoV-2/human/GBR/liverpool_strain/2020 (Genbank accession number MW041156.1, (*16*) and SARS-CoV-2 VOCs, Gamma (hCoV-19/England/520336_B1_P0/2021, GISAID ID: EPI_ISL_2080492); Delta (hCoV-19/England/BRS-UoB-233/2021, GISAID ID: EPI_ISL_15250227 (*17*) Omicron BA.1 (hCoV-19/England/BRS-UoB-942/2022, GISAID ID: EPI_ISL_18150332) and Omicron BA.4 (hCoV-19/England/BRS-UoB-942/2022, GISAID ID: EPI_ISL_18151340). The Delta, Omicron BA.1 and BA.4 VOCs were isolated from clinical research samples collected through the Bristol Biobank (UK NHS Research Ethics Committee Ref 20/WA/0273). The Gamma VOC was kindly provided by Professor Wendy Barclay, Imperial College, London and Professor Maria Zambon, UK Health Security Agency). Viral stocks were produced by infecting VeroE6-TMPRSS2 cells at a MOI of 0.01 in infection medium (Minimal Essential Medium (MEM) supplemented with 2 % FBS, L-glutamine, PenStrep and NEAAs. Virus containing supernatant was collected at 24-72 h post infection and clarified either by centrifugation for 10 min at 300 x g or filtration through a 0.2 µM membrane, stored at −80°C and used for infections. Tick-borne encephalitis virus (TBEV) strain Kuutsalo-14_Ixodes_ricinus_Finland-2017 (GenBank MG589938.1) was produced by infecting SK-N-SH cells at MOI (multiplicity of infection) 0.001 in infection medium (DMEM, 2 % FBS, L-glutamine, PenStrep, 0.35 µM rapamycin (SelleckChem, S1039)). Virus-containing supernatant was collected at 72 h.p.i., pre-cleared and stored as described above. Virus titres were determined using plaque assay or TCID_50_ titrations in VeroE6-TMPRSS (CoV2) or SK-N-SH (TBEV) cells. SFV-ZsG, RSV-GFP, and VSV-GFP were produced in BHK-21, Hela Hep2, and Vero-E6 cells respectively. Cells were infected at an MOI of 0.1 for 22 h (SFV-ZsG and VSV-GFP) or 48 h (RSV-GFP), in MEM supplemented with 0.2% BSA, 2 mM L-glutamine, penstrep and NEAA. The collected media were centrifuged twice at 300xg for 10 min, at 4°C and the supernatants aliquoted and stored at −80°C.

DENV2 (ST) was a clinical isolate obtained from Singapore General Hospital while DENV1-2402, DENV3-863, and DENV4-2270 were clinical isolates obtained from the Early Dengue Infection and Outcome study (*34*).

### Infection inhibition assays

#### Pre-incubation of mɑ-DGN with virus

For infections with SARS-CoV-2 Finland/1/2020, RSV, SFV, TBEV, VSV and hAd5, cells were seeded in a ViewPlate-96 (Perkin Elmer, FIN.) at a concentration of 10,000 cells per well one day prior to the assay. The virus was mixed with 10 µM *m*ɑ-DGN or buffer control, incubated for 5 min at room temperature and used to infect the cells. For each virus, the viral dose was calibrated to obtain 20-30% infected cells in buffer control treated cells. Following a 16 h (TBEV, RSV-GFP, hAdV5) or 6 -hour (SFV-ZsG, VSV-GFP) infection, the cells were fixed with 4 % formaldehyde, washed with PBS and infected cells were identified by immunostaining using antibodies raised against dsRNA (SCICONS) for TBEV, or by direct detection of fluorescence produced by reporter fluorescent proteins expressed by recombinant viruses (SFV-ZsG, VSV-GFP, RSV-GFP, hAdV5-RFP), and quantified using the open source Cell Profiler-4 software. The inhibitory effect of ɑ-DGN was determined by comparing the percentage of infected cells in the presence of the protein or buffer control.

#### Pre-incubation of mɑ-DGN with cells

For infections with SARS-CoV-2 Wuhan, Gamma, VOC Delta Clin94, Omicron BA.1 and Omicron BA.4, and influenza A virus (X31) cells were seeded in clear 96-well Microplates (Greiner Bio-one, UK) at a concentration of 10,000 cells per well one day prior to the assay. Cells were pre-treated with 10 µM *m*ɑ-DGN for 30 min at 37°C prior to the addition of virus in infection media (MEM, 2 % FBS, 0.1 % NEAA) at a MOI of 0.1. Following a 16-hour incubation at 37°C and 5 % CO_2_, cells were fixed with 4 % formaldehyde. The fixed cells were blocked in PBS, 1 % BSA for 30 min followed by permeabilization for 3 min (PBS, 1 % BSA, 0.1 % Triton-X 100). Thereafter the cells were stained with anti-N (1:2,000) (SARS-CoV-2) or with anti-NP (1:15) (IAV) for 60 min at room temperature. The cells were washed and stained with goat anti-rabbit Alexa Fluor-488 (1:2,500, SARS-CoV-2) or goat anti-mouse Alexa Fluor-488 (1:2,500, IAV) for 30 min at room temperature. Hoechst was used to stain the nuclei. Cells were imaged using an automated spinning-disc microscope CQ1 (Confocal Quantitative Image 130 Cytometer, Yokogawa, Japan). Images were analysed using cell Path Finder (Yokogawa, Japan) software and graphed in Prism 9.4 (GraphPad), The inhibitory effect of ɑ-DGN was determined by comparing the percentage of infected cells in the presence or absence of the protein.

### Generation and viral infection of organoids

Human jejunal samples were obtained from patients undergoing Roux-en-Y gastric bypass surgery. The study regarding relevant samples and associated ethical regulations were approved by Helsinki University Hospital review board. Written and informed consent was obtained before enrolment. Crypts from human jejunal biopsies were isolated by vigorous shaking after 1 h incubation in ice cold PBS with 10 mM EDTA. To enrich crypts, tissue suspension was filtered through 70 μm nylon mesh. Enriched crypts were washed once with cold PBS and plated into 60 % Matrigel (BD Biosciences). overlaid with hENR medium; (Advanced DMEM/F12 (Gibco), 1× Glutamax (Gibco), 10 mM Hepes (AdDF++), 1× B-27 (Gibco), 1× N-2 (Gibco), 50 ng/ml of mouse EGF (RnD), 100 ng/ml noggin (Peprotech), 500 ng/ml of R-spondin-1 (RnD), 10 nM gastrin (Sigma-Aldrich), 100 ng/ml Wnt3A (RnD), 10 mM nicotinamide (Sigma-Aldrich), 500 nM A-83-01 (Sigma-Aldrich) and 10 μM SB202190 (Sigma-Aldrich). Organoids were collected into cold adDF++ medium and washed once to remove excess Matrigel. Organoids were then sheared using a P1000 pipet tip and trituration and onto multi-well tissue culture plates at approximately 500,000 cells per well. SARS-CoV-2 was mixed with 1 or 10 µM *m*ɑ-DGN or buffer, incubated for 5 min at room temperature and used to infect the broken organoids at MOI=1 for 2 h at 37°C and 5 % CO2, after which the unbound virus was removed by washing with AdDF++ and centrifugation for 5 min at 300g. Organoids were embedded in Matrigel incubated for 20 min at 37°C and 5 % CO2 to allow matrigel solidification and then overlayed with hENR. After 48 h organoids were fixed with 4 % formaldehyde and processed for immunofluorescence using an antibody against dsRNA.

### Pseudovirus particle generation and infection

Pseudovirus particles expressing the surface spike protein of interest (SARS-CoV-S, SARS-CoV-2-S, MERS-CoV-S, HCoV-229E-S) were generated using the VSVΔG system as previously described (*35, 36*). In Brief, HEK293T cells were transfected with pCG1_SARS-CoV-S, pCG1_SARS-CoV-2-S, pCAGGS_MERS-CoV-S or pCAGGS_HCoV-229E-S using Polyethylenimin (PEI) Max (MW = 40,000KDa, Polysciences, DE) as transfection reagent at a ratio of 4:1 (PEI:DNA) in serum free DMEM for 4 h at 37°C. Subsequently the cells were washed with PBS and cultured in DMEM supplemented with 5 % FBS at 37°C overnight. The cells were then subjected to the replication deficient VSV*ΔG-fLuc vector, whose glycoprotein (G) gene is substituted by an expression cassette for enhanced green fluorescent protein (eGFP) and firefly luciferase for 2 h at 37°C. The cells were washed once with PBS and cultured in medium supplemented with anti-VSV-G I1 antibody (Clone 8G5F11, Absolute Antibody, UK) for a further 24 h at 37°C. The supernatant was harvested and concentrated by ultracentrifugation through a sucrose cushion (10 %-30 %) at 100,000 x g for 2 h. The pellet was resuspended in PBS supplemented with 10 % FBS, aliquoted and stored at −80°C until use.

For inhibition studies using VSV pseudotyped particles (VSVpp) expressing different spike proteins (VSVpp-SARS-CoV-S, VSVpp-SARS-CoV-2-S, VSVpp-MERS-CoV-S and VSVpp-HCoV-229E-S), Caco-2 cells were grown in clear 96-well Microplates (Greiner Bio-one, UK). Cells were pre-treated with the respective compounds (*m*ɑ-DGN, *h*ɑ-DGN and *m*S6) for 30 min at 37°C prior to the addition of VSVpp in infection media (DMEM, 2 % FBS). Following a 16 hours incubation at 37°C, cells were fixed with 4 % formaldehyde, washed and Hoechst was used to stain for the nuclei. Infection was quantified by measuring GFP-positive cells using an automated spinning-disc microscope CQ1 (Confocal Quantitative Image 130 Cytometer, Yokogawa, Japan). Images were analysed using cell Path Finder (Yokogawa, Japan), graphed in Prism 9.4 (GraphPad), and IC50 values were calculated by nonlinear regression analysis using the dose–response (variable slope) equation.

### Plaque assays

Cells were grown in a 6-well plate and infected with 200 µl of 10-fold serial dilutions of virus in infection medium. Following 1 h incubation at 37°C and 5 % CO_2_ with rocking, cell monolayers were covered with 3 ml of overlay medium (MEM, 2 % FBS, L-glutamine, penstrep, 1.2 % Avicel). Cells were fixed with 10 % formaldehyde at 60 (CoV2) or 72 (TBEV) h.p.i. and plaques were visualized by staining with a crystal violet solution (0.2 % crystal violet, 1 % methanol, 20 % ethanol, 3.6 % formaldehyde). Viral titres were determined as plaque-forming units per ml of stock.

### mα-DGN binding to cells

For these experiments *m*α-DGN was conjugated with Alexa Fluor-647 succinimidyl (NHS) esther (ThermoFisher Scientific) following an established protocol (*37*). A549-ACE2-TMPRSS2 or Caco-2-ACE2 cells were seeded into clear 96-well Microplates (Greiner Bio-one, UK) at a concentration of 6,000 or 10,000 cells per well respectively the day before the binding assay. The cells were washed once with media and then incubated with 4 µM of Alexa Fluor-647 labelled *m*α-DGN for 45 min on a chilled metal plate on ice, after which cells were immediately fixed with 4 % formaldehyde. Fixed cells were then blocked with PBS, 1 % BSA for 30 min at room temperature. The nucleus was stained with Hoechst and phalloidin Alexa Fluor-594 (1:2,000) was used to stain for actin filaments. The cells were imaged using an automated spinning-disc microscope CQ1 (Confocal Quantitative Image 130 Cytometer, Yokogawa, Japan) with a 40x/0.95na objective. Images were generated using ImageJ.

### SARS-CoV-2 replicon transfection and inhibition assay

The Wuhan-Hu-1 SARS-CoV-2 (pSC2-Rep-Wu-Gp-RL) replicon, lacking the coding regions for the Spike and membrane proteins was used. These regions were replaced with coding regions for an enhanced GFP-puromycin N-acetyl transferase fusion protein and Renilla luciferase, respectively (*17*).

The transfection and inhibition assay were performed following previously published methods (*17*). Briefly, VTN cells were co-transfected using a Neon Transfection System (Invitrogen, ThermoFisher, UK) with 250 ng N-gene mRNA and 1 μg replicon genomic RNA. Immediately after transfection 10,000 cells were seeded into flat bottomed 96-well tissue culture plates or clear 96-well Microplates (Greiner Bio-one, UK) containing 10 μM *m*α-DGN, vehicle control or 1μM remdesivir in growth media. The plates were incubated at 37°C, 5% CO_2_ for 18 h.

Luciferase assay: To measure the Renilla luciferase activity, the Renilla Luciferase Assay System (Promega, UK) was used according to manufacturer’s instructions. Briefly, the cells from the flat bottomed 96-well tissue culture plates were carefully washed 1x with PBS and lysed with 1x Renilla Luciferase lysis buffer and stored at −20°C. To perform the luciferase assay the lysate was transferred into white LUMITRAC plates (Greiner Bio-one, UK) and Renilla luciferase activity was measured using a GloMAX® Explorer microplate reader (Promega, UK).

Immunofluorescence assay: The cells in the clear 96-well Microplates (Greiner Bio-one, UK) were fixed with 4% paraformaldehyde, blocked in PBS, 1 % BSA and permeabilized for 3 min (PBS, 1 % BSA, 0.1 % Triton-X 100). The cells were stained against double stranded-RNA (1:250), washed and stained with goat anti-mouse Alexa Fluor-647 (1:2,500) for 30 min at room temperature. Hoechst was used to stain the nuclei. Cells were imaged using an automated spinning-disc microscope CQ1 (Confocal Quantitative Image 130 Cytometer, Yokogawa, Japan) using a 10x objective. Images were analysed using cell Path Finder (Yokogawa, Japan) software and graphed in Prism 9.4 (GraphPad). The inhibitory effect of ɑ-DGN was determined by comparing the percentage of infected cells in the presence or absence of the protein.

### SARS-CoV-2 binding inhibition assay

HeLa cells stably expressing ACE2 were seeded into clear 96-well Microplates (Greiner Bio-one, UK) at the concentration of 6,000 cells per well the day before the assay. For the assay the cells were washed once with media and then pre-treated with 10 µM *m*α-DGN for 30 min at 37°C. Following the incubation, SARS-CoV-2 was added at an MOI=50 and allowed to bind on chilled metal plates for 45 min, after which cells were immediately fixed with 4 % formaldehyde. Fixed cells were then blocked with PBS, 1 % BSA for 30 min at room temperature. Cells were stained with a rabbit anti-S (1:200) antibody for 60 min at room temperature, washed and stained with a goat anti-rabbit Alexa Fluor-594 (1:2,500) secondary antibody for 30 min at room temperature. The nucleus was stained with Hoechst and Wheat Germ Agglutinin (WGA) Alexa Fluor-647 (1:200) was used to stain the cell membrane. The cells were imaged using an automated spinning-disc microscope CQ1 (Confocal Quantitative Image 130 Cytometer, Yokogawa, Japan) with a 40x/0.95na objective. The images were analysed by a pipeline created in Cell Path Finder (Yokogawa, Japan). Briefly, SARS-CoV-2-S-AF594 labelled particles within the boundary of a segmented cell were counted to quantify the number of bound viral particles per cell. More than 700 cells (100-250 cells per well) were analysed for each condition.

### Infection inhibition assay using DENV

#### Infection inhibition in Huh7 cells

Huh7 cells were pre-treated with *m*α-DGN or an unrelated protein control for 2 h before each of the four DENVs were inoculated at MOI 1 and incubated for 2 h at 37°C. The virus inoculum was then removed, and the cells were maintained in DMEM containing 9% FBS for a further 5 days. Cells and supernatants were collected at the specified time point and stored at −80°C before DENV infection levels were measured by real time-PCR. RNA extraction of cell lysates and supernatant were done using the RNeasy Mini Kit and QIAamp Viral RNA Mini Kit, respectively (Qiagen) according to the manufacturer’s instructions. Next, cDNA synthesis was performed using the qScript cDNA Synthesis Kit (Quantas Biosciences). Viral replication was measured by RT-qPCR using the SYBR Green Supermix Kit (Roche) with primers for pan-serotype DENV and for the TATA box binding protein as a reference (*38*). All reactions were run on a Roche LightCycler 480, and data analysis was performed with LightCycler 480 software.

#### Infection inhibition in human primary CD14+ monocytes

Human CD14+ monocytes were harvested by negative selection from peripheral blood mononuclear cells (PBMCs) of healthy donors according to a protocol approved by the National University of Singapore’s Institutional Review Board (reference no. B-15-227) as described previously (*39*). Monocytes were pre-treated with *m*α-DGN or the protein control for 2 h. Following this incubation monocytes were infected with DENV4 at an MOI of 10 and incubated for 2 h at 37°C. The virus was then washed off and fresh *m*α-DGN or unrelated protein control was added. 48 h later the supernatant was harvested for the determination of DENV titres by plaque assay, as previously described (*39*).

### Animal experiments

K18-hACE2 C57/BL/6 mice were obtained from InVivos, Singapore. All animal procedures were performed under approved protocols by the Institutional Animal Care and Use Committee at Singapore Health Services (protocol no: 2020/SHS/1554) and were in accordance with guidelines provided by the National Advisory Committee for Laboratory Animal Research (NACLAR) in Singapore.

Treatment with *m*α-DGN, *h*α-DGN and buffer control, as well as the SARS-CoV-2 challenges were done via the intranasal (IN) route. For the IN administrations mice were slightly anaesthetised with Isoflurane. In the first instance *m*αDGN was administered at 7.5 µg/mouse or 0.75 µg/mouse via the IN route, immediately followed by IN administration of a lethal (5×10^4 TCID50 in 25 µl PBS) SARS-CoV-2 (brain derived SG12-B; (hCoV-19/Singapore/2/2020)) challenge dose. In the second experiment animals received 7.5 µg *m*αDGN or *h*αDGN/mouse 2 h prior to the SARS-CoV-2 (lung derived SG12-L; (hCoV-19/Singapore/2/2020)) IN challenge (2×10^4 PFU in 25 µl PBS) (*19*) and then 7.5 µg *m*αDGN or *h*αDGN /mouse once a day for the 3 consecutive days after the challenge. Mice were observed daily for weight loss, clinical signs of disease and death. Mice were sacrificed when they reached 20% loss in bodyweight or a clinical score of 10. At the endpoint lung and brain tissues were collected and analysed for viral loads by plaque assay or qRT-PCR that targeted the ORF1ab gene of SARS-CoV-2, as previously described (*19, 40*).

### Statistical analysis

Statistical analysis was performed using GraphPad Prism software 9.4.0. The statistical test used for each experiment is stated in the corresponding figure legend. Statistical significance is indicated by asterisks and defined as p*<0.05, p**<0.01, p***<0.001, p****<0.0001. All experiments were carried out with multiple independent biological replicates (n≥3). Main Fig. 3: n= number of animals per group. Error bars indicating either SD or SEM are mentioned in the respective figure legends.

## Acknowledgments

ADD is a member of the G2P-UK United Kingdom National Virology consortium funded by the Medical Research Council/UKRI that supplied SARS-CoV-2 variants. The authors wish to thank Drs. Anne Juuti and Kirsi Pietiläinen (University of Helsinki) for kindly providing human biopsy samples. MGB wishes to thank Emmaline Stotter (University of Bristol) for helping with protein preparation.

## Author contributions

Conceptualization: MGB, KK, AB, YY

Methodology: MGB, KK, ESG, MA, ME

Investigation: MGB, KK, ESG, MA, SA, SA, OV, PK, ME

Visualization: MGB, KK, MA, GB

Funding acquisition: MGB, KK, ADD, SJB, IC, GB, AB, YY

Project administration: MGB, KK, GB, AB, YY

Supervision: MGB, KK, ADD, SJB, EEO, GB, YY

Writing – original draft: MGB, KK, AB, YY

Writing – review & editing: MGB, KK, ESG, MA, ME, ADD, SJB, IC, EEO, GB, AB, YY

## Data and materials availability

All data are available in the main text or the supplementary materials. Materials related to this study may be provided by MGB and AB upon request and through a material transfer agreement (MTA) with the University of Bristol.

## Supplementary Materials

**Fig. S1.**
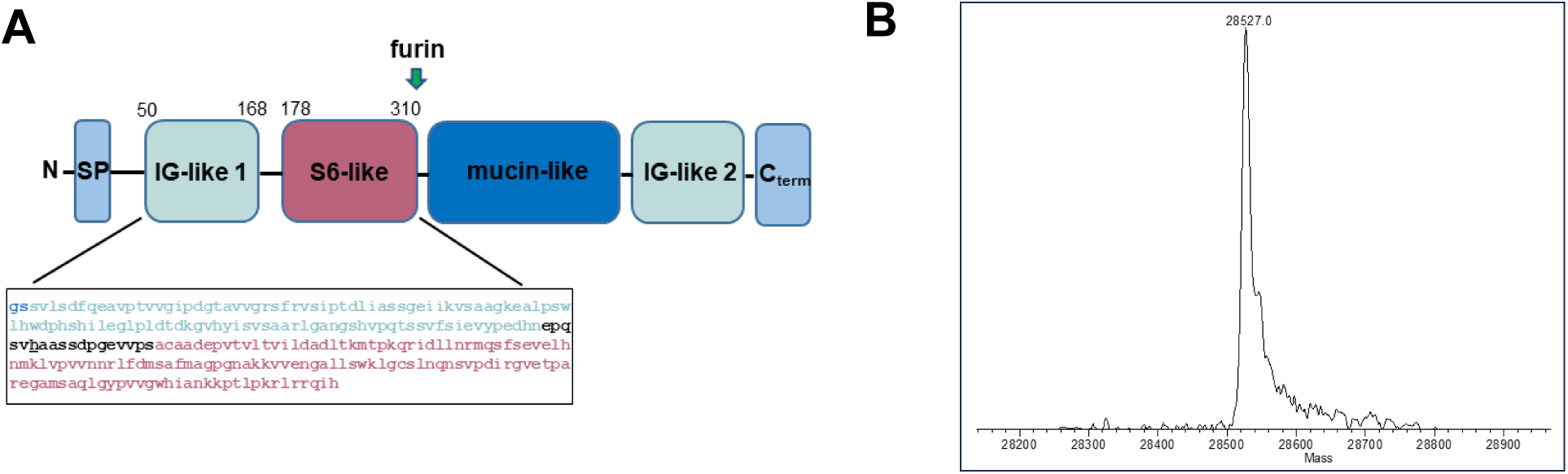
Recombinant α-DGN properties. (**A**) Domain structure of mouse α-DG (*m*α-DG). SP: signal peptide, Ig-like: immunoglobulin-like domain, S6: ribosomal protein S6-like domain, arrow: furin cleavage site, mucin-like: highly glycosylated central domain of α-DG, C-term: C-terminal portion. The amino acid sequence of the N-terminal domain (α-DGN) as cloned and expressed in this study, is indicated below the cartoon. The sequence of the Ig-like domain is indicated in cyan, the sequence of the S6-like domain is indicated in magenta. The His residue that replaced the original Arg166 in order to stabilize the recombinant product is in black underlined. (**B**) Electrospray Ionization Liquid Chromatography Mass Spectrometry profile of *m*α-DG. The recombinant product runs as a single, sharp peak with a MW of 28527Da, matching the MW calculated based on the amino acid sequence (28528Da).

**Fig. S2.**
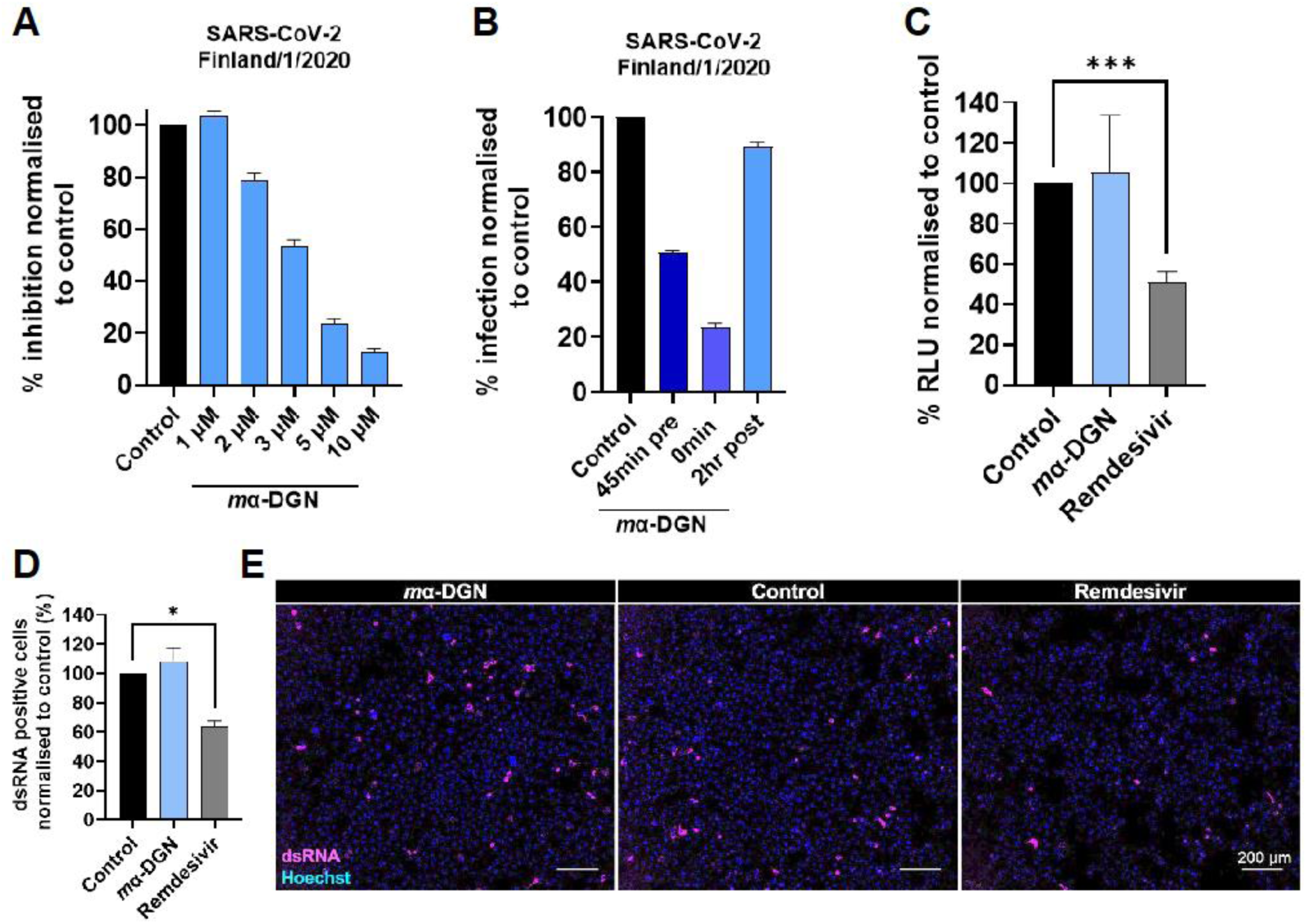
*m*α-DGN inhibitory activity against SARS-CoV2. (**A**) Dose dependent inhibitory activity of *m*α-DGN (1 µM-10 µM) against SARS-CoV-2 infection in HEK293T cells expressing human ACE2 and TMPRSS2. The graph shows the % inhibition of SARS-CoV-2 normalised to the buffer-treated control (means + SD, n=3). (**B**) Inhibitory activity of *m*α-DGN when pre-incubated with SARS-CoV-2 for 45 min, added at the same time as SARS-CoV-2 or added 2 h post infection with SARS-CoV-2. After a 16 h incubation infection was determined by viral N protein staining. Results are represented as % inhibition normalised to the control (means + SD). (**C**) Expression levels of Relative luminescence units (RLU) in SARS-CoV-2 replicon RNA transfected VTN cells treated for 18 h with 10 μM *m*α-DGN, vehicle control or 1 μM remdesivir. (**D**) VTN cells transfected with SARS-CoV-2 replicon RNA and treated with 10 μM mα-DGN, vehicle control or 1μM Remdesivir were fixed after 18 h incubation and stained for dsRNA. The graph shows the percentage of dsRNA positive cells after *m*α-DGN, or Remdesivir treatment normalized to the vehicle control treated cells. (**E**) Representative immunofluorescence images of dsRNA (magenta) positive cells after treatment with *m*α-DGN, vehicle control or Remdesivir. Scale bar: 200 µm.

**Fig. S3.**
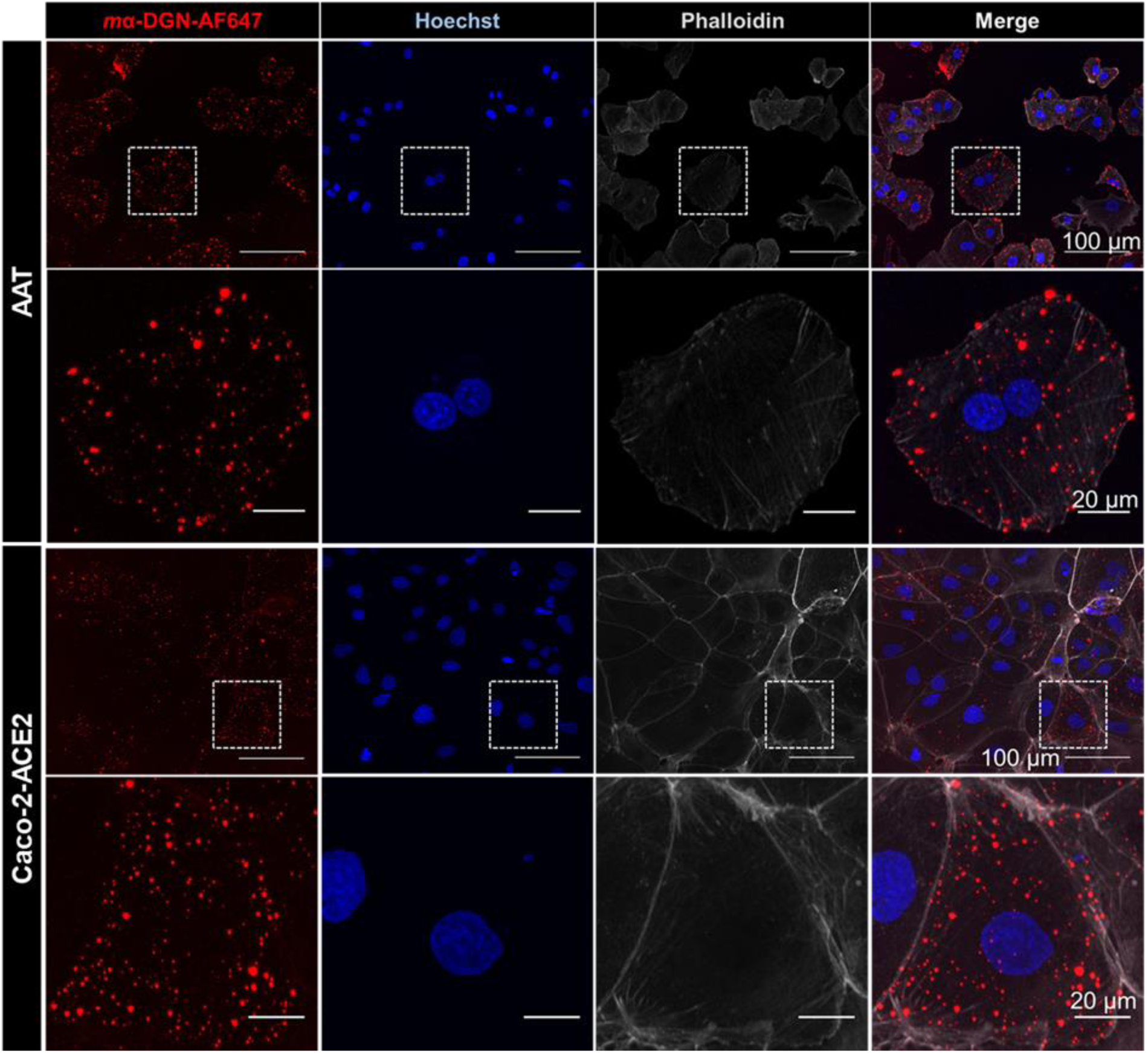
α-DGN binds to different cell lines. (**A**) A549-ACE2-TMPRSS2 (AAT) or (**B**) Caco-2-ACE2 cells were incubated with 4 µM Alexa Fluor-647 labelled *m*α-DGN for 45 min in the cold and fixed. Nuclei was stained with Hoechst (blue), actin with phalloidin-AF596 (grey) and *m*α-DGN Alexa Fluor-648 is shown in red. Scale bars, 100 µm and 20 µm in the magnified panels. In both panels, the highlighted square regions in the upper image rows were enlarged in the lower image rows.

**Fig. S4.**
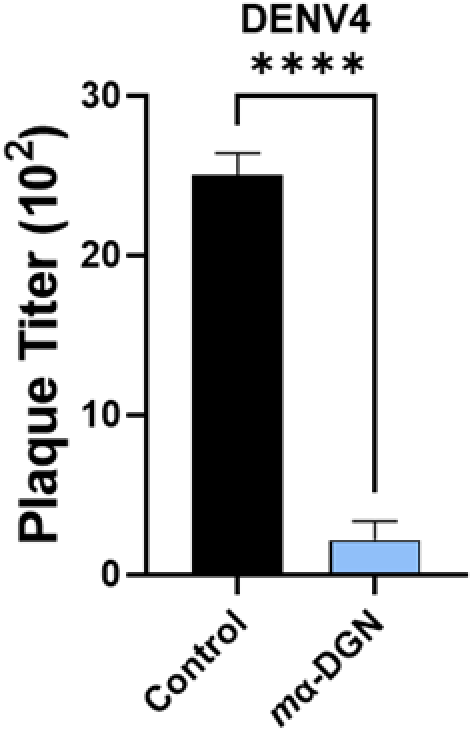
α-DGN inhibits DENV4 infection of human primary monocytes. (**A**) Human CD14+ monocytes were pre-treated with *m*α-DGN for 2 h prior to infection with DENV4 at an MOI of 10. Inoculum was washed off after 2 h and fresh *m*α-DGN added. After 48 h supernatant was harvested and the viral titre determined by plaque assay. Results are represented as plaque titres (10^2^) of n=6 replicates (means + SD).

## References

1. Barresi, R., and Campbell, K. P. (2006). Dystroglycan: from biosynthesis to pathogenesis of human disease. J. Cell. Sci. 119, 199–207. 10.1242/jcs.02814.

2. Sciandra, F., Bigotti, M. G., Giardina, B., Bozzi, M., and Brancaccio, A. (2015). Genetic engineering of dystroglycan in animal models of muscular dystrophy. Biomed. Res. Int. 2015, 635792. 10.1155/2015/635792.

3. Adams, J. C., and Brancaccio, A. (2015). The evolution of the dystroglycan complex, a major mediator of muscle integrity. Biol Open. 4, 1163–1179. 10.1242/bio.012468.

4. Cao, W., Henry, M. D., Borrow, P., Yamada, H., Elder, J. H., Ravkov, E. V., Nichol, S. T., Compans, R. W., Campbell, K. P., and Oldstone, M. B. (1998). Identification of α-dystroglycan as a receptor for lymphocytic choriomeningitis virus and Lassa fever virus. Science 282, 2079– 2081. 10.1126/science.282.5396.2079.

5. Katz, M., Weinstein, J., Eilon-Ashkenazy, M., Gehring, K., Cohen-Dvashi, H., Elad, N., Fleishman, S. J., and Diskin, R. (2022). Structure and receptor recognition by the Lassa virus spike complex. Nature 603, 174–179. 10.1038/s41586-022-04429-2.

6. Kanagawa, M., Saito, F., Kunz, S., Yoshida-Moriguchi, T., Barresi, R., Kobayashi, Y. M., Muschler, J., Dumanski, J. P., Michele, D. E., Oldstone, M. B., and Campbell, K. P. (2004). Molecular recognition by LARGE is essential for expression of functional dystroglycan. Cell 117, 953–964. 10.1016/j.cell.2004.06.003.

7. Brancaccio, A., Schulthess, T., Gesemann, M., and Engel, J. (1997). The N-terminal region of alpha-dystroglycan is an autonomous globular domain. Eur. J. Biochem. 246, 166–172. 10.1111/j.1432-1033.1997.00166.x.

8. Bozic, D., Sciandra, F., Lamba, D., and Brancaccio, A. (2004). The structure of the N-terminal region of murine skeletal muscle alpha-dystroglycan discloses a modular architecture. J. Biol. Chem. 279, 44812–44816. 10.1074/jbc.C400353200.

9. Okuma, H., Hord, J. M., Chandel, I., Venzke, D., Anderson, M. E., Walimbe, A. S., Joseph, S., Gastel, Z., Hara, Y., Saito, F., et al. (2023). N-terminal domain on dystroglycan enables LARGE1 to extend matriglycan on α-dystroglycan and prevents muscular dystrophy. eLife 12, e82811. 10.7554/eLife.82811.

10. Hesse, C., Johansson, I., Mattsson, N., Bremell, D., Andreasson, U., Halim, A., Anckarsäter, R., Blennow, K., Anckarsäter, H., Zetterberg, H., et al. (2011). The N-terminal domain of alpha-dystroglycan, released as a 38 kDa protein, is increased in cerebrospinal fluid in patients with Lyme neuroborreliosis. Biochem. Biophys. Res. Commun. 412, 494–499. 10.1016/j.bbrc.2011.07.129.

11. Saito, F., Saito-Arai, Y., Nakamura, A., Shimizu, T., and Matsumura, K. (2008). Processing and secretion of the N-terminal domain of alpha-dystroglycan in cell culture media. FEBS Lett. 582, 439–444. 10.1016/j.febslet.2008.01.006.

12. Crowe, K. E., Shao, G., Flanigan, K. M., and Martin, P. T. (2016). N-terminal α-dystroglycan (αDG-N): a potential serum biomarker for Duchenne muscular dystrophy. J. Neuromusc. Dis. 3, 247–260. 10.3233/JND-150127.

13. de Greef, J. C., Slutter, B., Anderson, M. E., Hamlyn, R., O’Campo Landa, R., McNutt, E. J., Hara, Y., Pewe, L. L., Venzke, D., Matsumura, K., et al. (2019). Protective role for the N-terminal domain of alpha-dystroglycan in Influenza A virus proliferation. Proc. Natl. Acad. Sci. U S A 116, 11396–11401. 10.1073/pnas.1904493116.

14. Sciandra, F., Schneider, M., Giardina, B., Baumgartner, S., Petrucci, T. C., and Brancaccio, A. (2001). Identification of the beta-dystroglycan binding epitope within the C-terminal region of alpha-dystroglycan. Eur. J. Biochem. 268, 4590–4597. 10.1046/j.1432-1327.2001.02386.x

15. Cantuti-Castelvetri, L., Ojha, R., Pedro, L. D., Djannatian, M., Franz, J., Kuivanen, S., van der Meer, F., Kallio, K., Kaya, T., Anastasina, M., et al. (2020). Neuropilin-1 facilitates SARS-CoV-2 cell entry and infectivity. Science 370, 856–860. 10.1126/science.abd2985.

16. Daly, J. L., Simonetti, B., Klein, K., Chen, K. E., Williamson, M. K., Anton-Plagaro, C., Shoemark, D. K., Simón-Gracia, L., Bauer, M., Hollandi, R., et al. (2020). Neuropilin-1 is a host factor for SARS-CoV-2 infection. Science 370, 861–865. 10.1126/science.abd3072.

17. Erdmann, M., Kavanagh Williamson, M., Jearanaiwitayakul, T., Bazire, J., Matthews, D. A., and Davidson, A. D. (2022). Development of SARS-CoV-2 replicons for the ancestral virus and variant of concern Delta for antiviral screening. bioRxiv, 10.1101/2022.10.11.511804.

18. Lamers, M. M., Beumer, J., van der Vaart, J., Knoops, K., Puschhof, J., Breugem, T. I., Ravelli, R. B. G., Paul van Schayck, J., Mykytyn, A. Z., Duimel, H. Q., et al. (2020). SARS-CoV-2 productively infects human gut enterocytes. Science 369, 50–54. 10.1126/science.abc1669.

19. Gan, E.S., Syenina, A., Linster, M., Ng, B., Zhang, S.L., Watanabe, S., Rajarethinam, R., Tan, H. C., Smith, G. J. and Ooi, E. E. (2021). A mouse model of lethal respiratory dysfunction for SARS-CoV-2 infection. Antiviral Res. 193, 105138. 10.1016/j.antiviral.2021.105138.

20. Pulkkinen, L. I. A., Barrass, S. V., Domanska, A., Överby, A. K., Anastasina, M., and Butcher, S.J. (2022). Molecular organisation of Tick-borne encephalitis virus. Viruses 14, 792. 10.3390/v14040792.

21. Spuul, P., Balistreri, G., Kääriäinen, L., and Ahola, T. (2010). Phosphatidylinositol 3-kinase-, actin-, and microtubule-dependent transport of Semliki Forest Virus replication complexes from the plasma membrane to modified lysosomes. J. virol. 84, 7543–7557. 10.1128/JVI.00477-10.

22. Krzyzaniak, M. A., Zumstein, M. T., Gerez, J. A., Picotti, P., and Helenius, A. (2013). Host cell entry of respiratory syncytial virus involves macropinocytosis followed by proteolytic activation of the F protein. PLoS pathog. 9, e1003309. 10.1371/journal.ppat.1003309.

23. Johannsdottir, H. K., Mancini, R., Kartenbeck, J., Amato, L., and Helenius, A. (2009). Host cell factors and functions involved in vesicular stomatitis virus entry. J. virol., 83, 440–453. 10.1128/JVI.01864-08.24.

24. Spiropoulou, C. F., Kunz, S., Rollin, P. E., Campbell, K. P., and Oldstone, M. B. (2002). New World arenavirus clade C, but not clade A and B viruses, utilizes alpha-dystroglycan as its major receptor. J. Virol. 76, 5140–5146. 10.1128/jvi.76.10.5140-5146.2002.

25. Kunz, S., Rojek, J. M., Kanagawa, M., Spiropoulou, C. F., Barresi, R., Campbell, K. P. and Oldstone, M. B. (2005). Posttranslational modification of alpha-dystroglycan, the cellular receptor for arenaviruses, by the glycosyltransferase LARGE is critical for virus binding. J. virol. 79, 14282–14296. 10.1128/JVI.79.22.14282-14296.2005.

26. Halstead, S. B. (2014). Dengue antibody-dependent enhancement: knowns and unknowns. Microbiol. Spectr. 2. 10.1128/microbiolspec.AID-0022-2014. 10.1128/microbiolspec.AID-0022-2014.

27. Sauna, Z. E., Lagassé, D., Pedras-Vasconcelos, J., Golding, B., and Rosenberg, A. S. (2018). Evaluating and mitigating the immunogenicity of therapeutic proteins. Trends biotechnol. 36, 1068–1084. 10.1016/j.tibtech.2018.05.008.

28. Widyasari, K., and Kim, J. (2023). A review of the currently available antibody therapy for the treatment of coronavirus disease 2019 (COVID-19). Antibodies (Basel) 12, 5. 10.3390/antib12010005. 10.3390/antib12010005.29.

29. Cho, J., Shin, Y., Yang, J. S., Kim, J. W., Kim, K. C., and Lee, J. Y. (2023). Evaluation of antiviral drugs against newly emerged SARS-CoV-2 Omicron subvariants. Antiviral Res. 214, 105609. 10.1016/j.antiviral.2023.105609.

30. Narayan, R., Sharma, M., Yadav, R., Biji, A., Khatun, O., Kaur, S., Kanojia, A., Joy, C. M., Rajmani, R., Sharma, P. R., et al. (2023). Picolinic acid is a broad-spectrum inhibitor of enveloped virus entry that restricts SARS-CoV-2 and influenza A virus in vivo. Cell Rep. Med. 4, 101127. 10.1016/j.xcrm.2023.101127.

31. Abdelnabi, R., Foo, C. S., Kaptein, S. J. F., Boudewijns, R., Vangeel, L., De Jonghe, S., Jochmans, D., Weynand, B., and Neyts, J. (2022). A SCID mouse model to evaluate the efficacy of antivirals against SARS-CoV-2 infection. J. Virol. 96, e0075822. 10.1128/jvi.00758-22.

32. Bozzi, M., Cassetta, A., Covaceuszach, S., Bigotti, M. G., Bannister, S., Hübner, W., Sciandra, F., Lamba, D., and Brancaccio, A. (2015). The structure of the T190M mutant of murine α-Dystroglycan at high resolution: insight into the molecular basis of a primary dystroglycanopathy. PLoS One 10, e0124277. 10.1371/journal.pone.0124277.

33. Liu, H., and Naismith, J. H. (2008). An efficient one-step site-directed deletion, insertion, single and multiple-site plasmid mutagenesis protocol. BMC biotechnol. 8, 91. 10.1186/1472-6750-8-91.

34. Low, J. G., Ooi, E. E., Tolfvenstam, T., Leo, Y. S., Hibberd, M. L., Ng, L. C., Lai, Y. L., Yap, G. S., Li, C. S., Vasudevan, S. G., and Ong, A. (2006). Early Dengue infection and outcome study (EDEN) - study design and preliminary findings. Ann. Acad. Med. Singap. 35, 783– 789.

35. Hoffmann, M., Kleine-Weber, H., and Pöhlmann, S. (2020). A multibasic cleavage site in the Spike protein of SARS-CoV-2 is essential for infection of human lung cells. Mol. Cell 78, 779–784.e5. 10.1016/j.molcel.2020.04.022.

36. Berger Rentsch, M., and Zimmer, G. (2011). A vesicular stomatitis virus replicon-based bioassay for the rapid and sensitive determination of multi-species type I interferon. PloS One 6, e25858. 10.1371/journal.pone.0025858.

37. Hayashi-Takanaka, Y., Stasevich, T. J., Kurumizaka, H., Nozaki, N., and Kimura, H. (2014). Evaluation of chemical fluorescent dyes as a protein conjugation partner for live cell imaging. PloS One 9, e106271. 10.1371/journal.pone.0106271.

38. Sun, B., Sundström, K. B., Chew, J. J., Bist, P., Gan, E. S., Tan, H. C., Goh, K. C., Chawla, T., Tang, C. K., and Ooi, E. E. (2017). Dengue virus activates cGAS through the release of mitochondrial DNA. Sci. Rep. 7, 3594. 10.1038/s41598-017-03932-139.

39. Chan, K. R., Zhang, S. L., Tan, H. C., Chan, Y. K., Chow, A., Lim, A. P., Vasudevan, S. G., Hanson, B. J., and Ooi, E. E. (2011). Ligation of Fc gamma receptor IIB inhibits antibody-dependent enhancement of dengue virus infection. Proc. Natl. Acad. Sci. U S A 108, 12479– 12484. 10.1073/pnas.1106568108.

40. Zhu, N., Zhang, D., Wang, W., Li, X., Yang, B., Song, J., Zhao, X., Huang, B., Shi, W., Lu, R. (2020). A novel coronavirus from patients with pneumonia in China, 2019. N. Engl. J. Med. 382, 727–733. 10.1056/NEJMoa2001017.

